# Connecting the Microenvironmental Niche to Treatment Response in Ovarian Cancer

**DOI:** 10.1101/452052

**Authors:** Maximilian Strobl, Matthew Wicker, Vikram Adhikarla, W. Andrew Shockey, Eszter Lakatos, Pantea Pooladvand, Ryan Schenk, Linggih Saputro, Chandler Gatenbee, Martijn Koppens, Salvador Cruz García, Robert Wenham, Mehdi Damaghi, Jill A. Gallaher

## Abstract

Ovarian cancer has the highest mortality rate of all gynecologic cancers, which may be attributed to an often late stage diagnosis, when the cancer is already metastatic, and rapid development of treatment resistance. We propose that the metastatic disease could be better characterized by observing interactions within the microenvironmental niche of the primary site that shapes the tumor’s early phenotypic progression. We present a mechanistic mathematical model of ovarian cancer that considers spatial interactions between tumor cells and several key stromal components. We demonstrate how spatial biomarker imaging data from the primary tumor can be analyzed to define a patient-specific microenvironment in the mathematical model. We then show preliminary results, using this model, that demonstrate how differences in the niche composition of a tumor affects phenotypic evolution and treatment response.

## I Introduction

Ovarian cancer detected before becoming metastatic has a 5-year survival rate of 93%. However, early detection before metastases occur is challenging for a variety of reasons; nevertheless, only 15% of cases are discovered in an early stage [1]. The majority of patients (85%) present with Stage 3 or 4 cancer and excessive metastases into the peritoneum, the lining of the cavity between the visceral organs and the abdominal wall [2], [3]. Intraperitoneal metastases are difficult to treat, and recurrence is frequent, with only ~29% survival at 5 years [1]. While many patients may have their cancers respond to initial and subsequent therapies, the tumor will eventually become treatment-resistant.

To better understand why resistance emerges it is crucial to elucidate the complex interplay between the tumor phenotypes and their microenvironment that drives tumor evolution. All ovarian cancers share a common growth pattern, in which the cancer first develops in the fallopian tube or on the ovaries. Subsequently it metastasizes into the peritoneal cavity either through the basement membrane or by exfoliation into the peritoneal fluid [2], [4]. Occasionally metastatic sites also form in more distant locations, such as the brain, liver, or lung [2], however this occurs in only ~15% of patients [5]. The cells forming the seeds for more distant metastases are derived either from the primary tumor, or from a secondary site in the peritoneum [6]. Thus, we hypothesize that the microenvironmental influences (i.e. niche) of the primary tumors drives the early evolution of ovarian tumor cells and can inform how cells might respond in metastatic sites when treatment is applied.

In the following, we present a mechanistic mathematical model, developed during the 7^th^ Integrated Mathematical Oncology (IMO) workshop, which aims to elucidate this interaction between the tumor, its microenvironment, and treatment. Our approach proposes to integrate patient-specific multiplexed imaging of biopsies, to delineate specific microenvironmental niches in the tumor, and mathematical modeling, to make predictions for tumor response based on those niches. We believe that through this combination of multiplexed imaging and mechanistic modelling it will be possible in the future to perform personalized *in silico* testing to delineate the drivers in a patient’s tumor ecosystem [7], and provide guidance and predictive tools for the clinic (e.g. [8]).

## II Vascular and immune considerations in ovarian cancer

Many stromal agents play a role in the development of ovarian cancer such as vasculature, collagen, fibroblasts, cytokines, and T-cell infiltration [2], [9], [10]. The role of all of these factors are challenging to experimentally dissect, as they form a complex, interlinked system (see [9], [11] for a review). For simplicity, we only outline what we assume to be the most important factors: the vasculature and the immune response. This will motivate the analysis and model of the tumor-stroma interactions at the primary site, which are discussed in the remainder of this report.

Despite the numerous stromal factors at play in cancer, a key factor with a critical role involves the creation and regulation of tumor vasculature. Ovarian cancers have the capacity to grow to more than 40 times the size of the ovary, finding the room to expand into the peritoneal cavity [12]. However, the diffusion limit of oxygen imposes strict size limits on avascular tumors, so persistent vessel recruitment (angiogenesis) is pivotal for continued tumor growth [13]. One of the key mechanisms used by tumors to induce angiogenesis is the production of a signaling molecule called Vascular Endothelial Growth Factor (VEGF), which also plays a significant role in physiologic angiogenesis of the female reproductive cycle [12]. As such, VEGF has been found to be an important predictive marker in various ovarian and uterine pathologies, including ovarian cancer, with high VEGF levels being associated with poorer survival rates [12], [14], [15].

The immune system also plays an important role in shaping the growth and evolutionary trajectory of the tumor. Cytotoxic T-cells are able to detect and remove aberrant cells, and have long been recognized as key effectors in tumor surveillance and prevention [16]. As such there are strong evolutionary forces acting on tumors to develop mechanisms to evade or manipulate the immune-surveillance [16]. Given the remarkable host of immune-evasion strategies displayed by ovarian tumors, this indicates that immune-pressure is likely to play a defining role in ovarian tumor evolution. One of the best understood mechanisms to overcome predation by cytotoxic T-cells is through the over-expression of programmed cell death ligand 1 (PD-L1) by the tumor. By interaction with its receptor on the T-cell, this ligand prevents the T-cell from killing the cancer cell, and can further result in deactivation of the T-cell [17]. The quantity and ratios of different immune cells and the expression of immune checkpoint regulatory ligands on tumors (e.g. PDL1 expression) have been observed to correlate with ovarian cancer virulence and prognosis [18], [19].

Both the vasculature and immune response largely influence and are influenced by ovarian cancer. However, the impact on disease progression and response to treatment is not just determined by summing up the impact of each. In addition to the spatial distribution of these factors, they can influence each other and the dynamics of the tumor in a nonlinear fashion. For example, the pro-angiogenic molecule VEGF has suppressor effects on the immune system [20]. To better understand the interactions and dynamics involved, we propose the following framework to quantify spatial distributions of cells within the primary ovarian cancer tissue and build a mathematical model that incorporates this spatial information to make predictions of cancer progression and treatment response.

## III Modeling Approach

In this section, we define the key interactions between tumor cells and their microenvironment, build a preliminary model to simulate these interactions, detail the methodology for defining the patient-specific microenvironmental niche from imaging data, and show some preliminary results on how different niches can affect tumor growth, evolution, and response to treatment.

### A Defining the Model Tumor-Stromal Interactions

To gain a first understanding of the influence that stromal factors have on tumor evolution in primary ovarian cancer, we used the simplified model of tumor promotive and suppressive actions of vasculature and immune-surveillance outlined in section II and depicted in the diagram in Figure 1. In normal tissue, blood vessels both supply nutrients that will allow cells to proliferate and are the conduits for release of the tumor suppressing immune cells. This already sets up a trade-off for optimal tumor growth in the tissue. But within this spatially and temporally variable environment, tumor cells could also evolve by changing their phenotype. Three critical aspects of cell behavior are modeled: i) its VEGF production, ii) its PD-L1 expression, and iii) its ability to survive low nutrient (oxygen, glucose, amino acids) conditions. These cell phenotypes were chosen because of their complex tumor-environmental feedback, which will be explained below.

**Figure 1.**
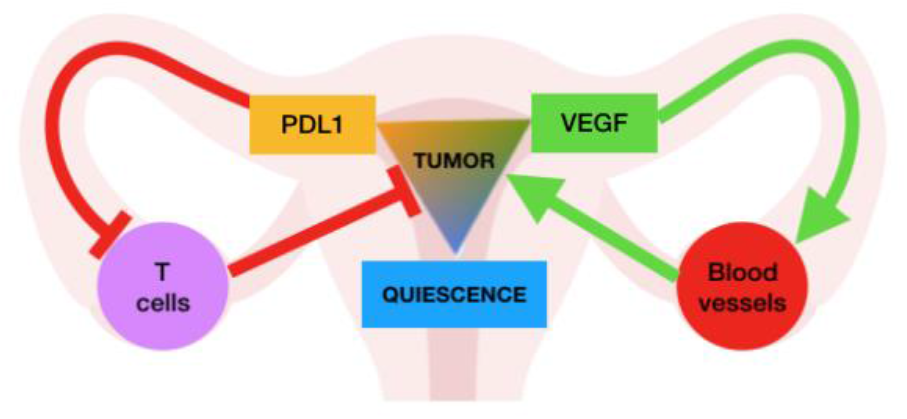
*Interactions between cancer cells and their environment in our model. Cancer cells use nutrients from blood vessels to grow and can be killed by T-cells, but they can also evolve adaptive traits. Cancer cells can secrete VEGF to recruit more blood vessels, express PD-L1 to avoid predation by T-cells and employ a quiescent strategy to avoid apoptosis in low nutrient areas*.

Cancer cells can recruit new vasculature by secreting VEGF. This niche construction by the tumor cells in turn increases the supply of nutrients, which promotes tumor growth. At the same time, vasculature releases T-cells, which predate on cancer cells. However, through the expression of PD-L1, a cancer cell can suppress the T-cell to avoid being killed. A third option for a cell to evolve is to lower the risk of extinction by becoming quiescent. By adopting a more dormant state the cell will not die as easily when nutrients are limited. It can therefore survive where tumor cells are dense and vasculature is low, but this comes at a cost of having a slow turnover even when nutrients are abundant. We then incorporate this system of interactions into a hybrid agent-based model.

### B A Hybrid Agent-Based Model

The model uses a 2-D hybrid cellular automata system implemented using the HAL framework in Java [21]. It contains both off-lattice agents (for cancer cells, T-cells, and vasculature) and continuous fields (for nutrient and chemotherapeutic drug concentrations). Simulations start with blood vessels and T-cells randomly distributed across the domain to define the stromal niche. A small tumor is initialized in the center of the domain. An example of the model setup is shown in Fig. 2.

**Figure 2.**
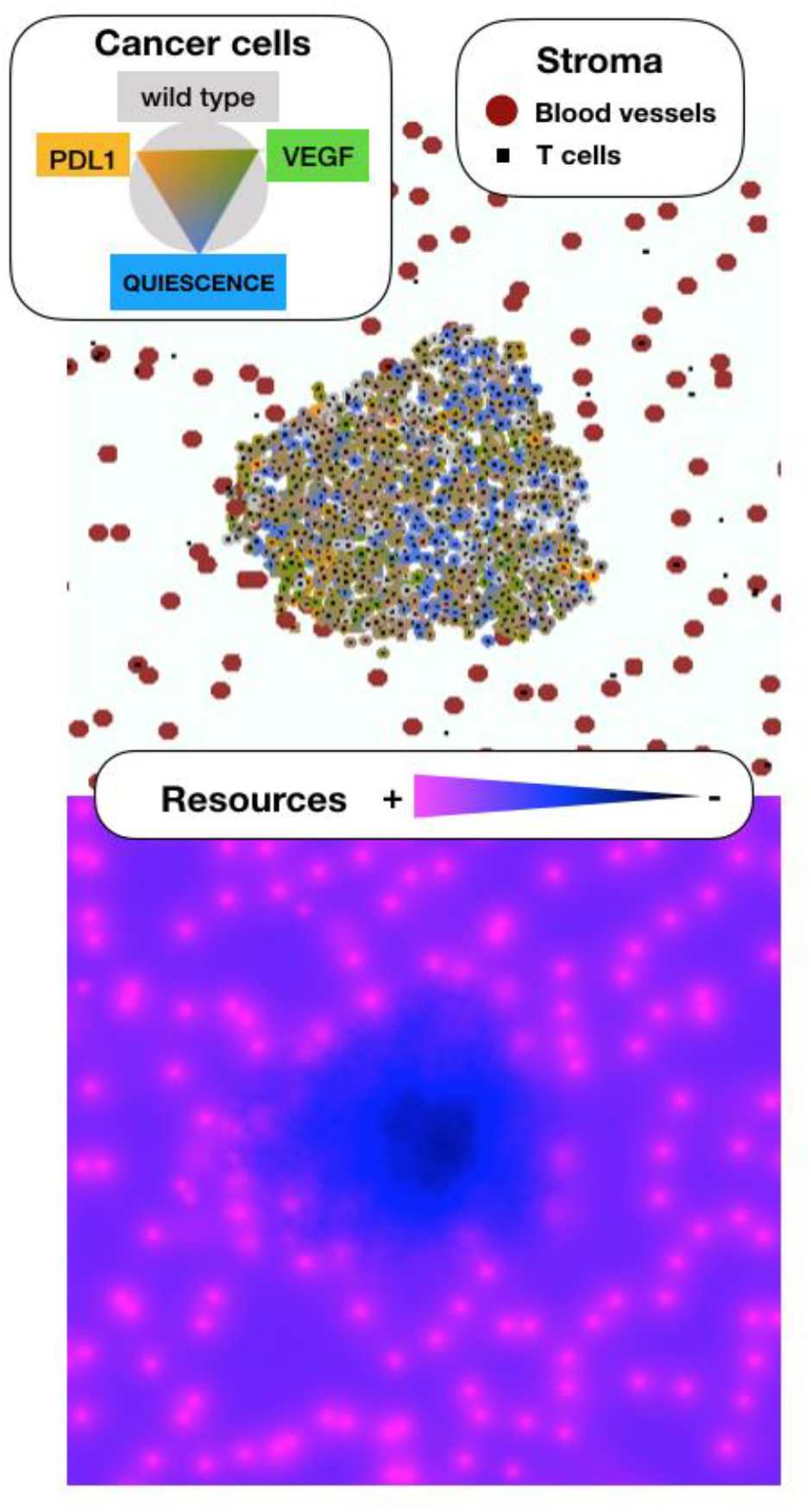
*Implementation of the modeling framework. There are three types of agents: cancer cells (colored by phenotype), blood vessels (red), and T-cells (green) shown in the top panel. Cancer cells can evolve to adopt different phenotypes, reflected by their color (see legend). The concentrations of nutrients supplied by the blood vessels is shown in the lower panel (pink: high, black: low)*.

In order to reduce the complexity of the model we represent the various nutrients used by cells (e.g. oxygen, glucose, amino acids) as a single resource. This resource is supplied by blood vessels and consumed by the cells at a fixed rate. Blood vessels in our model are represented by discrete agents and are dynamic in space and time. If the cell density around a vessel becomes too large, vessels will either be pushed into free space (if available) or collapse. New vessels are formed at a rate that increases with a higher local tumor burden and a higher expression of VEGF in the local vicinity. T-cells are also represented as individual agents. They originate from blood vessels and move in the domain in a Brownian fashion. If they are within a given radius of a cancer cell they will kill the cell with a probability that is inversely proportional to the level of PD-L1 expression. Reflecting the biology, T-cells can only kill a limited number of cancer cells before becoming exhausted and dying.

The third discrete agent in our model are the cancer cells. Depending on the amount of resources available in the environment and its phenotype, cancer cells will proliferate (high resource), quiesce (low resource), or die (very low resource). Each cancer cell is characterized by the following three traits: 1) the amount of VEGF it produces (promoting vasculature), 2) its PD-L1 expression (which lowers its probability of being killed by a T-cell), and 3) its quiescence range (the range of nutrient levels under which it is quiescent). Cells with greater quiescence range can survive under lower nutrient levels but also need higher nutrient levels to proliferate. Each trait is modelled as a value between 0 (no production/expression/interval width) and 1 (maximal production/expression/interval width). At each division, a mutation can occur with some probability *μ*, which will randomly perturb one of the three trait values.

Chemotherapy kills cells in the model proportional to the amount of drug at its location and the rate of cell proliferation. The concentration of nutrient and chemotherapy are modelled as continuous functions, whose spatiotemporal evolution is governed by partial differential equations (PDEs). The PDEs are solved numerically using a finite difference scheme, and the value experienced by each cell is interpolated from the computed solution using linear interpolation.

### C Defining the Stromal Niche

In order to gain insight into the type of environments that might be present in the primary tumor, we can use novel multiplexed imaging to generate spatial maps of different cellular or environmental features, as illustrated in Fig. 3. Shown is a 1mm punch biopsy from an epithelial ovarian tumor collected at the Moffitt Cancer Center. The tissue was imaged for 37 different cellular markers using imaging mass cytometry [22], performed by the Fluidigm Corporation (South San Francisco, CA, USA). Unfortunately, the set of markers did not include any of the key elements in our model. However, for illustration of the method, the set of panels in Fig. 3A shows the observed intensities of three different markers: E-cadherin – marker of epithelial cells, CD44 – marker of cell adhesion, and Collagen I – marker for most abundant collagen type. From this raw data it is possible to generate maps of the tumor environment, by segmenting the cells and analyzing their molecular expression profile. In the middle panel we show such a segmentation. The cells were segmented by first identifying their nuclei from the Histone stain, followed by estimating the cell outlines using Voronoi Tessellation (Fig. 3B). For more details on the individual cell phenotypes we take the mean level of each of the markers present within the cell’s area to identify tumor and stroma components of the tissue (Fig. 3C). The so obtained spatial distribution of different cell types can then be further analyzed using, for example, a Strauss point process model to identify and characterize environmental niches. We conclude that great detail on the composition of the tumor microenvironment can be obtained from multiplexed imaging. However, this information is static, and does not provide details about the underlying dynamics.

**Figure 3.**
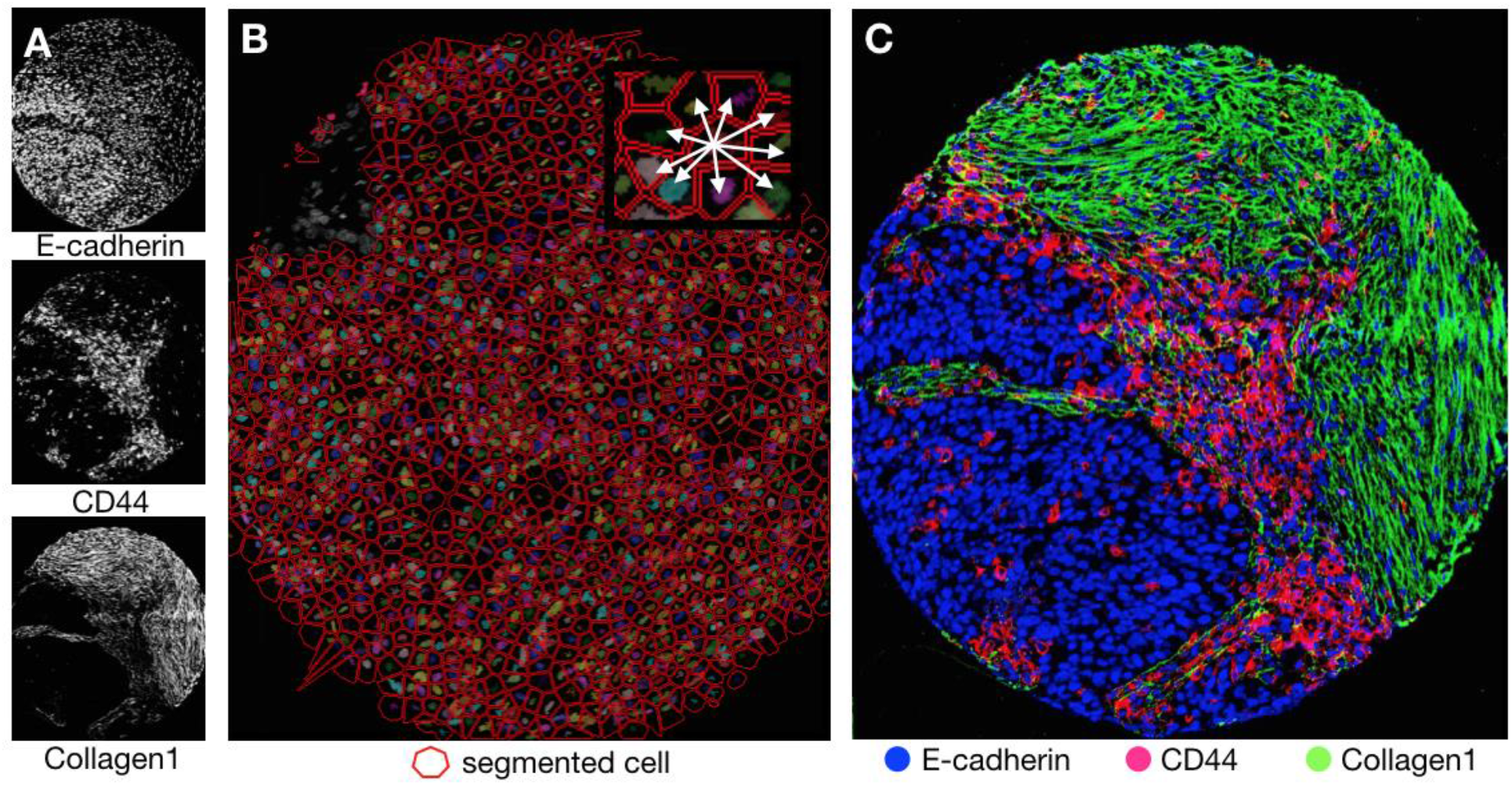
*Niche analysis using multiplexed imaging data. A) Imaging of separate biomarkers. B) Segmentation of cells. C) Composite of markers defines a stromal map that can parameterize and initialize the spatial mathematical model*.

This kind of patient-specific spatial map of a tumor’s niche can be obtained from imaging and used to parameterize and initialize the mathematical model. This would allow us to simulate the dynamics that could potentially arise from different treatments within a unique empirical spatial composition. Because we did not have data on the specific markers needed for this model, a couple of distinct microenvironmental niches were created to demonstrate how the tumor context can affect tumor growth and treatment response.

### D Preliminary Computational Results

Tumors were grown in 2 microenvironmental niches. Niche A was poorly vascularized but also had a low rate of immune infiltration whilst Niche B was highly vascularized but had a high rate of immune infiltration. For each case, the tumors were grown to 800 cells, at which point chemotherapy was applied for 10 days. This drug schedule was repeated every 50 days until reaching 350 days. The results are shown in Figure 4 for Niche A (Fig. 4A) and Niche B (Fig. 4B).

**Figure 4.**
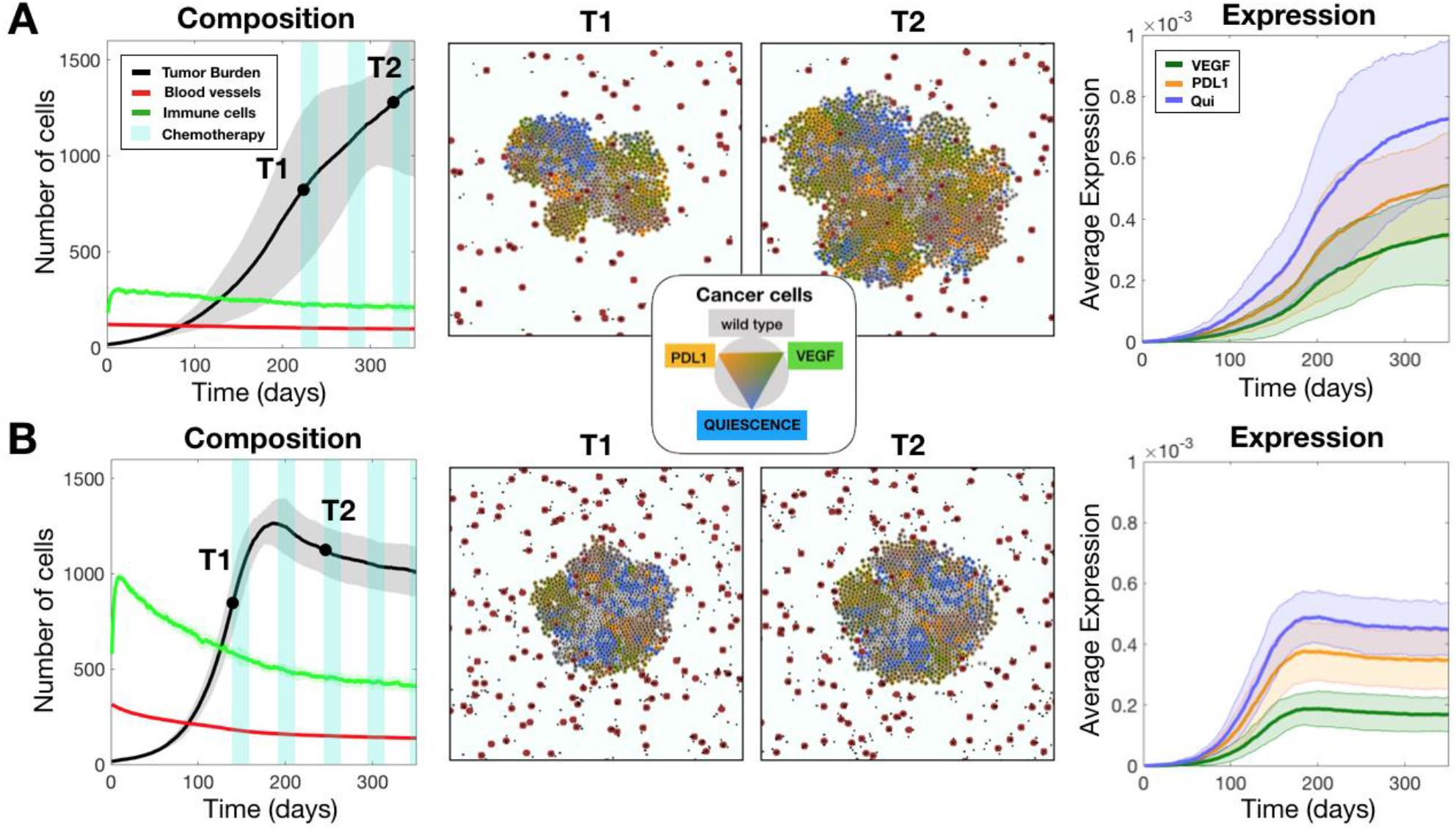
*Tumors were grown in two environments: A) a poorly vascularized niche with low immune infiltration and B) a well-vascularized niche with high immune infiltration. Each tumor is grown to 800 cells, and then chemotherapy is applied for 10 days at every 50 days interval. The leftmost graphs show the tissue composition in terms of the total number of tumor cells, the number of blood vessels, and the number of immune cells along with the applied chemotherapy schedule. The middle plots show the spatial compositions colored according to their relative expression level just prior to treatment initiation (T1), and then again after 2 cycles of treatment (T2). The rightmost plot shows the average expression levels for each of the 3 resistance mechanisms: VEGF, PDL1, and the quiescence phenotype. The error bars show the standard deviation from 10 runs*.

Niche A had very low resources, so the tumor grew slower than in Niche B where the high vascularization provided an abundance of resources and caused very rapid growth. The tumor in Niche B also grew more uniformly outward compared to the tumor in Niche A, which tended to bulge outward upon developing higher levels of expression of multiple phenotypes. However, once chemotherapy was applied, the slow growing tumor in Niche A did not show much response, while the faster growing tumor in Niche B decreased in size. This response might be due mostly to the difference in vascular density - more vessels delivered more drug to the tumor.

There were also differences in the tumor compositions in each niche. For both tumors, the highest expression level was the quiescence type, followed by PDL1, and then VEGF. However, there was a difference in the expression levels and their distributions amongst the cells. The low resource environment of Niche A selected for higher expression of all resistance mechanisms during growth with more variance than in Niche B. The tumor in Niche A took a longer time to reach the same size before chemotherapy is applied, so the proliferation rate, and therefore, also the rate of accumulating higher expression, was also low. However, combined with a lack of immune predation, Niche A allowed higher expression levels to accumulate, which slowed turnover and made the cells less responsive to chemotherapy. Perhaps a tumor in this niche would respond better to a more targeted therapeutic approach. Nevertheless, we did ignore the fact that cells evolved in the primary might behave differently when encountering a different microenvironment at the metastatic site and instead assumed that cells coming from a specific primary niche might seek out a comparable metastatic niche. This decoupling could be important and should be considered in future studies.

## IV Conclusion

In this report, we presented an approach to better understand the role of stroma in the evolution and treatment response of ovarian cancers. Tumor-stroma interactions, which are often neglected in models of cancer, can significantly shape the evolution of tumor cell phenotypes, and ultimately affect treatment outcomes. Here, we identified several key stromal influences on ovarian cancer and built a hybrid agent-based model to simulate tumor growth and chemotherapy treatment. We outlined methods for analyzing spatial data using mass cytometry data from ovarian cancer tissue samples to investigate the presence of different niches in the primary tumor. Given an appropriate set of stromal markers, the method presented here could be extended to provide an in-depth analysis of spatial interactions and allow for patient specific tests of treatment plans.

The time frame of the workshop was too short for large scale data analysis or integration of the data into the model, but we were able to explore with the model how 2 different niches affected tumor growth, composition, and response to treatment. We found that a high density of blood vessels and strength of immune-surveillance caused faster growth before treatment, but a larger population decline during treatment application compared to the niche with a low density of blood vessels and low immune-surveillance. Furthermore, we found that the low resource environment selected for higher expression levels of all phenotypes. Our simulations confirm that niche can drive phenotypic evolution and therefore affect tumor composition and treatment response. Overall, we hope that this report illustrates that the spatial structure of non-tumor components has great potential to affect tumor composition. We envision that future computational models will incorporate more patient-specific stromal interactions guided by multiplexed imaging data to improve understanding of ovarian cancer and personalize treatment response predictions.

## About the Authors

M. S. is with the Wolfson Centre for Mathematical Biology, University of Oxford, Oxford, UK.

M. W. is at the Department of Computer Science, University of Georgia, GA, USA.

V. A. is in the Division of Mathematical Oncology at City of Hope National Medical Center, Duarte, CA, USA.

A. S. is with the Department of Biomedical Engineering at Georgia Institute of Technology and Emory University, Atlanta, GA, USA.

E. L. is with the Barts Cancer Institute, Queen Mary University of London, London, UK.

P. P. is at the School of Mathematics and Statistics, University of Sidney, Sidney, Australia.

R. S. and M. K. are with the Wellcome Centre for Human Genetics, University of Oxford, Oxford, UK.

L. S. is at the Department of Bioinformatics and Computational Biology at University of South Florida, Tampa, FL, USA.

S. C.-G. is with Escuela Superior de Apan, Universidad Autónoma del Estado de Hidalgo, Apan, Hidalgo, Mexico.

M. D., J. G., C. G., R.S. and R. W. are with the H. Lee Moffitt Cancer Center & Research Institute, Tampa, FL, USA.

*,^+^Equal contributions

## Acknowledgments

We would like to thank the IMO Chair, Dr. Alexander Anderson, for organizing the 7^th^ Annual Moffitt IMO workshop: Stroma, where this project was conceived. We are also extremely grateful to the Moffitt Cancer Center and the Moffitt PSOC for supporting this workshop through the NCI U54CA193489 grant. In addition, we would like to thank Anthony Magliocco and Douglas Marchion for making the multiplexed imaging data available to us.

## References

[1] C. K. (eds). Howlader N, Noone AM, Krapcho M, Miller D, Bishop K, Kosary CL, Yu M, Ruhl J, Tatalovich Z, Mariotto A, Lewis DR, Chen HS, Feuer EJ, “Contents of the SEER Cancer Statistics Review (CSR), 1975-2014,” SEER Cancer Stat. Rev. 1975-2014, Natl. Cancer Institute., vol. 2015, pp. 31–32, 2016.

[2] J. O. A. M. van Baal et al., “Development of Peritoneal Carcinomatosis in Epithelial Ovarian Cancer: A Review,” J. Histochem. Cytochem., p. 002215541774289, 2017.

[3] E. Lengyel, “Ovarian cancer development and metastasis,” Am. J. Pathol., vol. 177, no. 3 pp. 1053–1064, 2010.

[4] J. A. Gubbels, N. Claussen, A. K. Kapur, J. P. Connor, and M. S. Patankar, “The detection, treatment, and biology of epithelial ovarian cancer,” J. Ovarian Res., vol. 3, no. 1, pp. 1–11, 2010.

[5] W. R. Robinson, J. Beyer, S. Griffin, and P. Kanjanavaikoon, “Extraperitoneal metastases from recurrent ovarian cancer,” Int. J. Gynecol. Cancer, vol. 22, no. 1 pp. 43–46, 2012.

[6] T.-L. Yeung, C. S. Leung, K.-P. Yip, C. L. Au Yeung, S. T. C. Wong., and S. C. Mok, “Cellular and molecular processes in ovarian cancer metastasis. A Review in the Theme: Cell and Molecular Processes in Cancer Metastasis,” Am. J. Physiol. - Cell Physiol., vol. 309, no. 7 pp. C444–C456, 2015.

[7] A. R. A. Anderson and V. Quaranta, “Integrative mathematical oncology,” Nat. Rev. Cancer, vol. 8, no. 3 pp. 227–234, 2008.

[8] J. Zhang, J. J. Cunningham, J. S. Brown, and R. A. Gatenby, “Integrating evolutionary dynamics into treatment of metastatic castrate-resistant prostate cancer,” Nat. Commun., vol. 8, no. 1 pp. 1–9, 2017.

[9] M. P. Steinkamp et al., “Ovarian Tumor Attachment, Invasion, and Vascularization Reflect Unique Microenvironments in the Peritoneum: Insights from Xenograft and Mathematical Models,” Front. Oncol., vol. 3, no. May, pp. 1–18, 2013.

[10] B. Davidson, C. G. Trope, and R. Reich, “The Role of the Tumor Stroma in Ovarian Cancer,” Front. Oncol., vol. 4, no. May, pp. 1–11, 2014.

[11] N. Weidner, “Intratumor microvessel density as a prognostic factor in cancer,” Am. J. Pathol., vol. 147, no. 1 pp. 9–19, 1995.

[12] D. A. Hazelton and T. C. Hamilton, “Vascular endothelial growth factor in ovarian cancer.,” Curr. Oncol. Rep., vol. 1, no. 1 pp. 59–63, 1999.

[13] M. A. Gimbrone, S. B. Leapman, R. S. Cotran, and J. Folkman, “Tumor Dormancy In Viva by Prevention of Neovascularization,” J. Exp. Med., vol. 136, pp. 261–276, 1972.

[14] W. Graybill, A. K. Sood, B. J. Monk, and R. L. Coleman, “State of the science: Emerging therapeutic strategies for targeting angiogenesis in ovarian cancer,” Gynecol. Oncol., vol. 138, no. 2 pp. 223–226, 2015.

[15] T. J. Duncan et al., “Vascular Endothelial Growth Factor Expression in Ovarian Cancer: A Model for Targeted Use of Novel Therapies?,” Clin. Cancer Res., vol. 14, no. 10 pp. 3030–3035, 2008.

[16] T. B. Turner, D. J. Buchsbaum, J. M. Straughn, T. D. Randall, and R. C. Arend, “Ovarian cancer and the immune system — The role of targeted therapies,” Gynecol. Oncol., vol. 142, no. 2 pp. 349–356, 2016.

[17] E. J. Wherry, “T cell exhaustion,” Nat. Immunol., vol. 12, no. 6 pp. 492–499, 2011.

[18] J. Hamanishi et al., “Programmed cell death 1 ligand 1 and tumor-infiltrating CD8+ T lymphocytes are prognostic factors of human ovarian cancer.,” Proc. Natl. Acad. Sci. U. S. A., vol. 104, no. 9 pp. 3360–5, 2007.

[19] S. Aust et al., “Absence of PD-L1 on tumor cells is associated with reduced MHC i expression and PD-L1 expression increases in recurrent serous ovarian cancer,” Sci. Rep., vol. 7, no. January, pp. 1–12, 2017.

[20] T. Voron et al., “Control of the Immune Response by Pro-Angiogenic Factors,” Front. Oncol., vol. 4, no. April, pp. 1–9, 2014.

[21] R. Bravo, M. Robertson-Tessi, and A. R. A. Anderson, “Hybrid Automata Library,” bioRxiv Prepr., pp. 1–24, 2018.

[22] C. Giesen et al., “Highly multiplexed imaging of tumor tissues with subcellular resolution by mass cytometry,” Nat. Methods, vol. 11, no. 4 pp. 417–422, 2014.

